# Radiolabeling Isolated Mitochondria with Tc-99m: A First-in-Field Protocol and Early Feasibility Findings

**DOI:** 10.1101/2025.06.15.658690

**Authors:** Melanie Walker, Francisco Javier Miralles, Keiko Prijoles, Jacob S. Kazmi, Jennifer Hough, David Lewis, Michael R. Levitt, Yasemin Sancak

**Affiliations:** Department of Neurological Surgery, University of Washington, Seattle, WA, USA; Stroke & Applied Neurosciences Center, University of Washington, Seattle, WA, USA; Department of Neurology, University of Washington School of Medicine, Seattle, WA, USA; Department of Pharmacology, University of Washington, Seattle, WA, USA; Donald and Barbara Zucker School of Medicine at Hofstra/Northwell, Hempstead, NY, USA; Laboratory for Critical Care Physiology, Feinstein Institutes for Medical Research, Manhasset, NY, USA; Division of Nuclear Medicine, University of Washington, Seattle, WA, USA; Department of Radiology, University of Washington School of Medicine, Seattle, WA, USA; Department of Mechanical Engineering, University of Washington School of Medicine, Seattle, WA, USA

**Author notes:** **Corresponding Author:** Melanie Walker, MD; 325 9^th^ Avenue, Harborview Medical Center, Department of Neurological Surgery, Seattle, WA USA.

**Keywords:** Electron Microscopy, Mitochondria, Mitochondrial Transplantation, Radiopharmaceuticals, SPECT, Technetium-99m

## Abstract

Mitochondrial transplantation is a promising but still experimental strategy for treating ischemic and metabolic disorders. A key barrier to its advancement is the lack of scalable, non-invasive methods for tracking transplanted extracellular mitochondria *in vivo*. Technetium-99m (Tc-99m) radiopharmaceuticals, widely used in SPECT imaging, may offer a clinically compatible solution. Cryopreserved mitochondria derived from HEK-293 cells were incubated with Tc-99m sestamibi, tetrofosmin, pertechnetate, or control solutions. After brief incubation and washing, mitochondrial pellets were analyzed for retained radioactivity. ATP content was measured to assess metabolic function, and electron microscopy was used to evaluate ultrastructural integrity. Tc-99m sestamibi and tetrofosmin showed labeling efficiencies of 2.74% and 2.68%, respectively. Pertechnetate demonstrated minimal uptake (0.34%). Radiolabeled mitochondria retained ATP production comparable to controls. Electron microscopy showed preserved double membranes and cristae. Controls confirmed assay specificity and viability. To our knowledge, this is the first report of radiolabeling isolated mitochondria with clinically approved Tc-99m agents. This platform supports the development of SPECT-compatible protocols for visualizing viable transplanted mitochondria in recipient tissues.

## INTRODUCTION

Mitochondrial transplantation is an emerging experimental strategy in regenerative medicine, particularly for ischemic stroke and other diseases associated with impaired cellular energetics and ischemia-reperfusion injury.^1,2^ Preclinical studies suggest that isolated mitochondria can be taken up by host cells and may help restore mitochondrial function and tissue bioenergetics.^2,3^ Although early human studies have demonstrated procedural safety,^4,5^ clinical development remains limited by practical barriers, including the absence of scalable and non-invasive methods to assess transplanted mitochondrial location, activity and viability after delivery.

Technetium-99m (Tc-99m) is one of the most widely used diagnostic radioisotopes in nuclear medicine, with favorable characteristics including a 140 keV gamma emission suitable for SPECT imaging, a six-hour half-life, and high availability from generator systems.^6^ Lipophilic, cationic Tc-99m radiopharmaceuticals such as sestamibi and tetrofosmin are routinely used in cardiac perfusion imaging and are known to accumulate in mitochondria based on membrane potential-dependent uptake.^7,8^ These agents demonstrate stable retention in the inner mitochondrial membrane, making them promising candidates for mitochondrial-specific labeling.

While the mitochondrial uptake mechanisms of Tc-99m agents are well described in intact cells and tissues,^7–9^ their behavior in isolated mitochondria has not been systematically studied. A method to do so could support non-invasive tracking of transplanted mitochondria *in vivo*, a key unmet need in the advancement of mitochondrial therapeutics.

Here, we present a first-in-field protocol for radiolabeling isolated extracellular mitochondria with Tc-99m sestamibi, tetrofosmin, and pertechnetate. We assess labeling efficiency, ATP retention, and ultrastructural preservation under mild conditions using unmodified mitochondria and commercially available reagents.

## MATERIALS AND METHODS

### Radiation Safety

All radiopharmaceutical handling and experimental procedures were conducted in accordance with federal regulations under U.S. NRC 10 CFR Part 35, Washington State Department of Health radiation protection guidelines, and University of Washington Radiation Use Authorization Permit R-05131, as reviewed by the institutional Radiation Safety Committee. Radiopharmaceutical handling took place in a dedicated, shielded hot lab within the institutional nuclear medicine facilities. The laboratory was equipped with lead-lined workstations, contamination monitoring systems, and controlled access, in compliance with institutional and federal radiation safety protocols. All personnel involved in radiolabeling procedures were certified in radiation safety and wore personal dosimeters throughout the study.

### Mitochondrial Preparation

Cryopreserved mitochondria (MRC-Q, LUCA Science, Tokyo, Japan), derived from human embryonic kidney (HEK-293) cells, were provided under Material Transfer Agreement #57121A and used without modification. Vials were removed from –80°C storage and immediately placed in a 25°C water bath for 2 minutes, with the cap held above the water line to prevent contamination. The outer surface was wiped dry, and the vial was transferred to an ice bucket. Contents were gently mixed by pipetting under a biosafety cabinet and maintained on ice throughout the labeling workflow, per the manufacturer’s instructions. Protein concentration was measured using a Bradford assay (Bio-Rad, Hercules, CA, USA), and aliquots were standardized to 50 µg total protein in 0.5 mL respiration buffer per condition. This minimal-handling approach was selected to evaluate labeling feasibility under conditions consistent with preclinical and translational workflows.^5,10^

### Radiopharmaceutical Preparation and Labeling

Tc-99m sestamibi (Cardiolite®, Lantheus), tetrofosmin (Myoview™, GE Healthcare), and sodium pertechnetate (Drytec™, GE Healthcare) were obtained from the institutional nuclear pharmacy (Cardinal Health, Seattle, WA) and prepared on the day of use according to manufacturer protocols. Each agent was diluted in 100 µL of sterile saline and then further diluted in 2.4 mL of respiration buffer to yield 2.5 mL total volume per condition. The buffer^11^ contained 250 mM sucrose, 2 mM KH_2_PO_4_, 10 mM MgCl_2_, 20 mM K-HEPES (pH 7.2), and 0.5 mM K-EGTA (pH 8.0). To promote oxidative phosphorylation and preserve mitochondrial membrane potential during labeling, the buffer was freshly supplemented with 5 mM glutamate and 5 mM malate prior to radiopharmaceutical addition.^12^ All components were prepared using molecular biology-grade reagents and sterile-filtered before use.

For each labeling condition, 50 µg of mitochondria (from previously quantified aliquots, as described above) were suspended in a final volume of 0.5 mL and incubated with 200 µCi of Tc-99m radiopharmaceutical at room temperature for 5 minutes. Gentle agitation was applied throughout the incubation to minimize mitochondrial aggregation or shear-related membrane damage. Two control conditions were included for each agent: One containing mitochondria in buffer without radiopharmaceutical (unlabeled control), and one consisting of buffer only (no mitochondria or radiopharmaceutical) to assess nonspecific background signal. Following incubation, samples were centrifuged at 9,000 × g for 5 minutes at 4°C. The supernatant was carefully aspirated, and the residual mitochondrial pellet was washed in ice-cold buffer to remove unbound radiotracer.

### Radioactivity Measurement

Retained radioactivity was quantified immediately following the wash step using a CRC-25R radionuclide activity calibrator (Capintec, NJ, USA), configured for Tc-99m detection at 140 keV. Mitochondrial pellets were transferred using low-retention pipette tips into pre-labeled 5 mL polypropylene tubes, which were placed at a fixed geometry within the calibrator to ensure measurement consistency. Calibration of the instrument was verified using a Tc-99m reference source before each measurement session, and background activity was recorded and subtracted. All measurements were corrected for radioactive decay based on the time elapsed from radiopharmaceutical preparation to sample readout. To maintain accuracy and linearity, calibrator performance was verified in accordance with established best practices for small-scale radiopharmaceutical preparation^13^. Data were recorded in microcuries (µCi) with a resolution of ±0.01 µCi. Each condition was measured in triplicate.

### Post-Labeling Metabolic Activity

To evaluate preservation of mitochondrial function following radiolabeling, ATP levels were measured in labeled and control pellets after freeze–thaw. Mitochondrial samples were resuspended in 10 µL of respiration buffer, snap-frozen on dry ice, and stored at −80°C for 24 hours. Samples were then thawed on ice and processed using the ATPLite Luminescence Assay System (Revvity, formerly PerkinElmer; Cat. No. 6016941), which detects ATP via luciferase-catalyzed bioluminescence. The assay was performed according to the manufacturer’s protocol. Luminescence was recorded in relative light units (RLU) using a Synergy H1 microplate reader (BioTek Instruments, Winooski, VT, USA) with a 10-second integration time. Samples were read in 96-well format using two technical replicates per condition. Buffer-only and unlabeled mitochondrial controls were included in each run to verify assay specificity and establish baseline signal.

### Transmission Electron Microscopy

Ultrastructural integrity of mitochondria was evaluated by transmission electron microscopy (TEM) following radiolabeling. Only tetrofosmin-labeled mitochondria were processed for TEM, alongside unlabeled controls. Mitochondrial pellets were fixed in 2.5% glutaraldehyde in 0.1 M sodium cacodylate buffer (pH 7.4) for 1 hour at 4°C, rinsed in the same buffer, and post-fixed in 1% osmium tetroxide for 1 hour at room temperature. Samples were dehydrated through a graded ethanol series and embedded in epoxy resin. Ultrathin sections (70–90 nm) were prepared using a diamond knife on an ultramicrotome (Leica Microsystems) and mounted on copper grids. Sections were stained with 2% uranyl acetate followed by lead citrate. Imaging was performed using a Tecnai G2 transmission electron microscope (Thermo Fisher Scientific) operated at 80–120 kV. Digital images were acquired at magnifications ranging from 20,000× to 100,000×. Morphometric scoring and blinded image analysis were not performed; imaging was intended for gross structural assessment.

### Quantification of Radiolabeling Efficiency

Labeling efficiency was defined as the percentage of administered Tc-99m activity retained in the mitochondrial pellet following incubation and a single wash, calculated as:

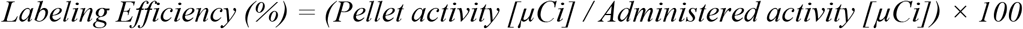

This approach is consistent with established protocols for radiolabeling human cells, including red blood cells,^14^ platelets,^15^ and leukocytes,^16^ as well as the standardized methodologies outlined in IAEA guidance.^17^ Administered activity was recorded at the time of tracer addition and decay-corrected using the Tc-99m half-life (6.01 hours, λ = 0.1151 hr^−1^). Post-wash activity in mitochondrial pellets was measured immediately to minimize decay-associated error. All measurements were performed using the consistent geometry and container format described above, with background subtraction applied to each sample, following operational guidance for radiopharmacy measurement reproducibility^.18^

The study was conducted under small-scale, research-use-only conditions. Per-sample activity levels (∼0.2 mCi) reflected common practice in feasibility-phase radiolabeling protocols and remained below thresholds requiring full GMP compliance.^13^ Process qualification and analytical control at this scale are consistent with current guidance on early-phase radiopharmaceutical validation^.19^

### Data Analysis and Statistical Reporting

All measurements were performed in triplicate unless otherwise stated. Data was recorded in Microsoft Excel and analyzed using GraphPad Prism (v9.0). Results are reported as mean ± standard deviation (SD). Inferential statistics were not applied, consistent with the exploratory and descriptive scope of this feasibility study.

## RESULTS

### Radiolabeling Efficiency

Among the tested Tc-99m agents, sestamibi and tetrofosmin demonstrated measurable uptake by isolated mitochondria, with mean labeling efficiencies of 2.74% and 2.68%, respectively. Pertechnetate showed lower labeling efficiency of 0.34%. These values represent the percentage of total administered radioactivity retained in the mitochondrial pellet following brief incubation and a single wash. Buffer-only and unlabeled mitochondria controls exhibited no measurable retained radioactivity, supporting specificity of uptake. Replicate data are shown in Table 1.

**Table 1.** Labeling Efficiency of Tc-99m Radiopharmaceuticals in Extracellular Mitochondria.

| Condition | Mean % (SD) | Calculated Dose ($\mu\text{Ci}$) | Actual Dose ($\mu\text{Ci}$) | Measured Radioactivity in Pellet ($\mu\text{Ci}$) | Labeling Efficiency (%) |
| --- | --- | --- | --- | --- | --- |
| **Sestamibi** | 2.74 (0.42) |  |  |  |  |
| Replicate 1 |  | 200 | 144.6 | 4 | 2.77 |
| Replicate 2 |  | 200 | 144.6 | 4 | 2.77 |
| Replicate 3 |  | 200 | 144.6 | 4.7 | 3.25 |
| **Tetrofosmin** | 2.68 (1.36) |  |  |  |  |
| Replicate 1 |  | 200 | 168.2 | 2.1 | 1.25 |
| Replicate 2 |  | 200 | 168.2 | 2.6 | 1.55 |
| Replicate 3 |  | 200 | 168.2 | 5.36 | 3.19 |
| **Pertechnetate** | 0.34 (0.24) |  |  |  |  |
| Replicate 1 |  | 200 | 183 | 0.18 | 0.1 |
| Replicate 2 |  | 200 | 183 | 0.73 | 0.4 |
| Replicate 3 |  | 200 | 183 | 1.3 | 0.71 |

### Post-Labeling ATP Production

Radiolabeled mitochondria exhibited detectable ATP levels following snap-freezing and thawing (Figure 1). ATP measurements in labeled samples were comparable to those in unlabeled mitochondria. Buffer-only samples showed no detectable ATP, confirming assay specificity.

**Figure 1.**
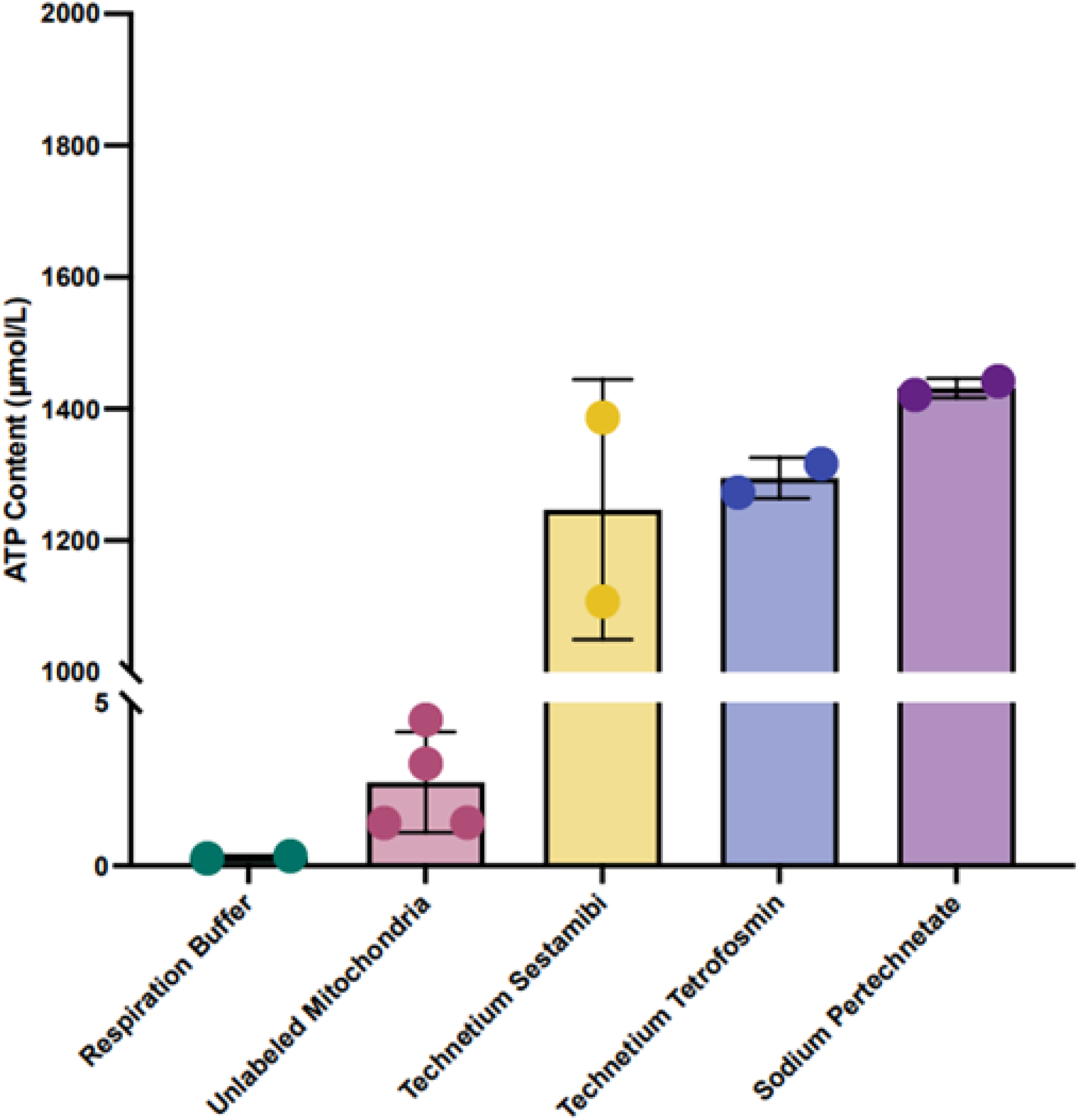
ATP Levels in Mitochondria Following Tc-99m Radiolabeling. Radiolabeled mitochondrial pellets (sestamibi, tetrofosmin, pertechnetate) exhibited detectable ATP content at a single post-labeling timepoint. Buffer-only controls showed no measurable ATP signal. Values reflect raw luminescence and were not normalized beyond total protein input (50 µg per replicate). *n* = 1 per group except for unlabeled mitochondria (*n* = 2); two technical replicates were performed per sample.

### Ultrastructural Integrity by Electron Microscopy

Tetrofosmin-labeled mitochondria were selected for ultrastructural analysis as the best-performing and most consistent tracer among those tested, with sufficient retained activity and sample availability for fixation. TEM revealed preserved gross ultrastructural features, including intact outer and inner membranes and well-defined cristae. No quantitative morphometric analysis or blinded scoring was performed; imaging was used solely to assess qualitative preservation of structure. Representative images are shown in Figure 2.

**Figure 2.**
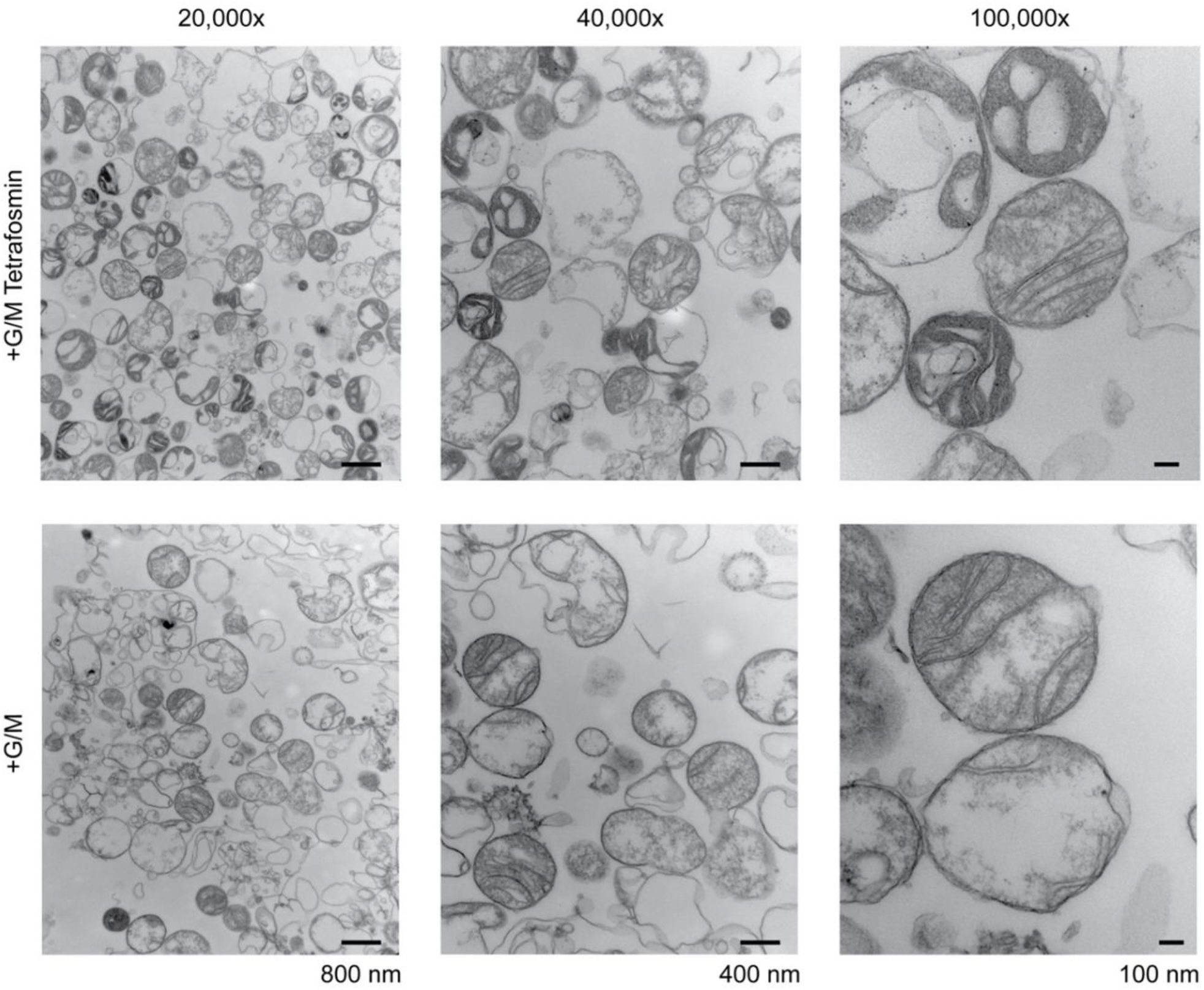
TEM of Mitochondria Radiolabeled with Tc-99m Tetrofosmin. Representative TEM images of mitochondrial pellets labeled with tetrofosmin, alongside unlabeled controls. Gross structural features, including intact double membranes and visible cristae, were qualitatively preserved. No morphometric analysis or blinded scoring was performed. Scale bars and magnification values are indicated on the figure panels. G/M indicates glutamate/malate supplementation.

## DISCUSSION

This study explored the preliminary feasibility of radiolabeling isolated mitochondria with Tc-99m agents commonly used in clinical nuclear imaging. While mitochondrial transplantation is under investigation as a therapeutic approach for ischemic and metabolic disorders,^1,4,5,10^ methods for non-invasively tracking extracellular mitochondria *in vivo* remain limited. However, such tracking is critical to evaluating the transport, uptake and eventual location of transplanted mitochondria. Preclinical models have demonstrated the uptake and functional rescue potential of transplanted mitochondria in injured tissues, with tracking often limited to fluorescence or luminescence-based techniques.^1,3,9^ These methods are poorly suited to clinical translation due to limited tissue penetration, photobleaching, and regulatory constraints.

Clinical radiolabeling protocols for cell-based products,^17^ including white blood cells,^16^ red blood cells,^14^ platelets,^15^ and stem cells, have been standardized over decades and are recognized by regulatory agencies.^17^ However, radiolabeling workflows established for cells have not been extended to subcellular organelles such as mitochondria, and no regulatory precedent currently exists. While clinical radiolabeling of organelles has not been reported, the physicochemical properties of Tc-99m agents support the scientific plausibility of this approach.^20–22^ Feasibility studies are needed to determine whether mitochondria can be efficiently and functionally radiolabeled with Tc-99m agents.

To our knowledge, the use of Tc-99m radiopharmaceuticals to label isolated, extracellular mitochondria has not previously been reported. Tc-99m radiopharmaceuticals such as sestamibi and tetrofosmin may offer a clinically scalable alternative due to their compatibility with SPECT-CT, short half-life, and regulatory precedent.^6–8^ Our results indicate that both sestamibi and tetrofosmin are capable of labeling isolated mitochondria to a modest degree, under minimally manipulated conditions. These agents are lipophilic cations with delocalized positive charge and moderate molecular weight, properties that allow passive diffusion across lipid bilayers and accumulation within polarized mitochondria via the negative inner membrane potential.^7,8,21^ Pertechnetate, which lacks lipophilicity and remains an anionic species in solution,^6,23^ showed markedly lower uptake. Sestamibi, an isonitrile compound, is more hydrophobic and typically exhibits slower washout and higher mitochondrial retention *in vivo*, whereas tetrofosmin, a diphosphine complex, is less lipophilic and displays faster blood clearance and lower membrane-binding affinity.^21,22,24,25^ These physicochemical differences may contribute to the variability observed between the two agents in labeling efficiency. Observed variability likely reflects the unoptimized, exploratory nature of the labeling protocol, including non-ideal incubation times, radiotracer concentrations, and post-labeling washes.

Radiolabeled mitochondria retained measurable ATP content after labeling and freeze-thaw, with levels similar to those in unlabeled controls. While slightly elevated ATP values were observed in some labeled samples, this are likely related to small sample size and potential variability in luminescence-based assays. One speculative explanation involves transient mitochondrial stimulation due to low-level ion flux or calcium mimicry by Tc-99m complexes, possibly mediated via the mitochondrial calcium uniporter.^26,27^ However, further validation is needed to determine membrane potential, oxygen consumption, and reactive oxygen species. Electron microscopy confirmed preservation of gross mitochondrial ultrastructure following tetrofosmin labeling, with clear visualization of intact double membranes and cristae. These findings support structural integrity under the labeling conditions, though further testing is required to confirm retained bioenergetic function.

Several limitations should be noted. Labeling conditions were not optimized for uptake kinetics, tracer retention, or radiotracer-to-mitochondria ratio. Free radiotracer adhesion to tubes and background signal were not independently quantified. Measurement variability may have been introduced by radioactive decay, sample geometry, or dose calibrator sensitivity. Sample sizes were small, and variability related to mitochondrial source or tracer handling may have contributed to signal differences. *In vivo* imaging, biodistribution, and clearance studies were not part of this early-stage effort.

These results demonstrate that clinically available Tc-99m agents can label isolated mitochondria at detectable levels without compromising ATP-generating capacity or gross membrane integrity. This study establishes a foundational protocol for adapting SPECT-compatible radiopharmaceuticals to the tracking of transplanted extracellular mitochondria. Building on this initial feasibility, ongoing work will focus on optimizing labeling conditions, applying orthogonal functional assays, and evaluating *in vivo* biodistribution and clearance in preclinical models.

## ACKNOWLEDGEMENTS

The authors thank the University of Washington Nuclear Medicine Pharmacy for assistance with radiopharmaceutical preparation, and the Electron Microscopy Core Facility for support with sample processing. We are also grateful to Ms. Ashtyn Winter (Stroke & Applied Neurosciences Center) for her contributions to mitochondrial acquisition and transport while employed as a research scientist, and to Ms. Joyce Chambers (Program Operations Specialist and Health Physicist, Environmental Health and Safety Radiation Safety) for her guidance on institutional radiation protocols. Mitochondrial organelle complex (MRC-Q) derived from human embryonic kidney 293 (HEK293) cells was provided by LUCA Science, Inc. under a materials transfer agreement.

## AUTHOR CONTRIBUTIONS

MW conceived the project, developed the radiolabeling protocol, and led manuscript preparation. YS, FJM, and MRL contributed to experimental design and critical review. KP, FJM, and JH performed sample handling and data acquisition. JK assisted with ATP quantification. DL and JH provided radiopharmaceutical access and imaging interpretation. All authors reviewed and approved the final manuscript.

## STATEMENTS AND DECLARATIONS

### Ethical approval

Not applicable. No human participants, animal models, or patient data were used in this study.

### Consent to participate

Not applicable.

### Consent for publication

Not applicable.

### Conflict of interest

MRL: Unrestricted educational grants from Medtronic and Stryker; consulting agreement with Aeaean Advisers, Metis Innovative, Genomadix, AIDoc and Arsenal Medical; equity interest in Proprio, Stroke Diagnostics, Apertur, Stereotaxis, Fluid Biomed, Synchron and Hyperion Surgical; editorial board of Journal of NeuroInterventional Surgery; Data safety monitoring board of Arsenal Medical. All other authors declare that they have no conflict of interest.

### Funding

This work received no external or internal funding. No financial support was provided by governmental, foundation, or commercial entities.

### Data availability

All data generated or analyzed during this study are contained in the manuscript.

